# Estimating Cardiac 5-HT2B Safety Margins for Repeated Low-Dose Psilocybin Using an Exposure–Response Model

**DOI:** 10.64898/2026.07.19.739440

**Authors:** William J. Tyler, Edward Sellers, Michael B. McDonnell

**Affiliations:** Diamond Therapeutics USA, Inc., Birmingham, Alabama, USA; Diamond Therapeutics, Inc., Toronto, Ontario, Canada

**Keywords:** psilocybin, psilocin, 5-HT2B receptor, valvular heart disease, partial agonism, exposure–response modeling, receptor pharmacology, drug safety

## Abstract

Repeated low-dose psilocybin is being developed as a scalable outpatient treatment for mood and anxiety disorders, but chronic exposure raises concern because psilocin binds the cardiac serotonin 5-HT2B receptor, whose sustained agonism causes drug-induced valvular heart disease (VHD). We evaluated this risk using an exposure–response model that incorporates functional efficacy and exposure duration rather than binding affinity alone. Plasma psilocin concentrations were converted into the time-integrated increment in 5-HT2B G_q_ signaling above endogenous serotonergic tone (ΔTIA) and calibrated against drugs and conditions with known valvular outcomes. All modeled exposures known to cause human VHD scored ΔTIA ≥ +172 %·h/day, whereas exposures not associated with VHD scored ≤ +28. A candidate 3 mg daily psilocybin regimen scored ΔTIA +3, roughly two orders of magnitude below the weakest valvulopathic exposure. This safety margin arises from psilocin’s low-efficacy partial agonism at 5-HT2B (E_max_ ≈51.8% of serotonin, compared with 96% for norfenfluramine) and its short half-life (≈ 2.5 h), which prevents accumulation and produces brief daily receptor engagement. In support of the model, rats receiving continuous psilocin for 12 days at plasma concentrations ≈2.4-fold above the projected human peak for 3 mg daily psilocybin showed no valvular lesions by blinded histopathology. This exposure duration however cannot exclude slowly developing fibrosis. Emerging human data, including serial echocardiography in repeated LSD microdosing and a large observational cohort, are also agreement with the model. Collectively, these findings suggest a favorable safety margin for daily, sub-hallucinogenic psilocybin use in clinical indications. Nevertheless, continued pharmacological and clinical investigations should include prospective echocardiographic monitoring to advance the clinical safety profile of sub-hallucinogenic psilocybin and support its evaluation across a broad array of therapeutic programs.

**Graphical Abstract:** 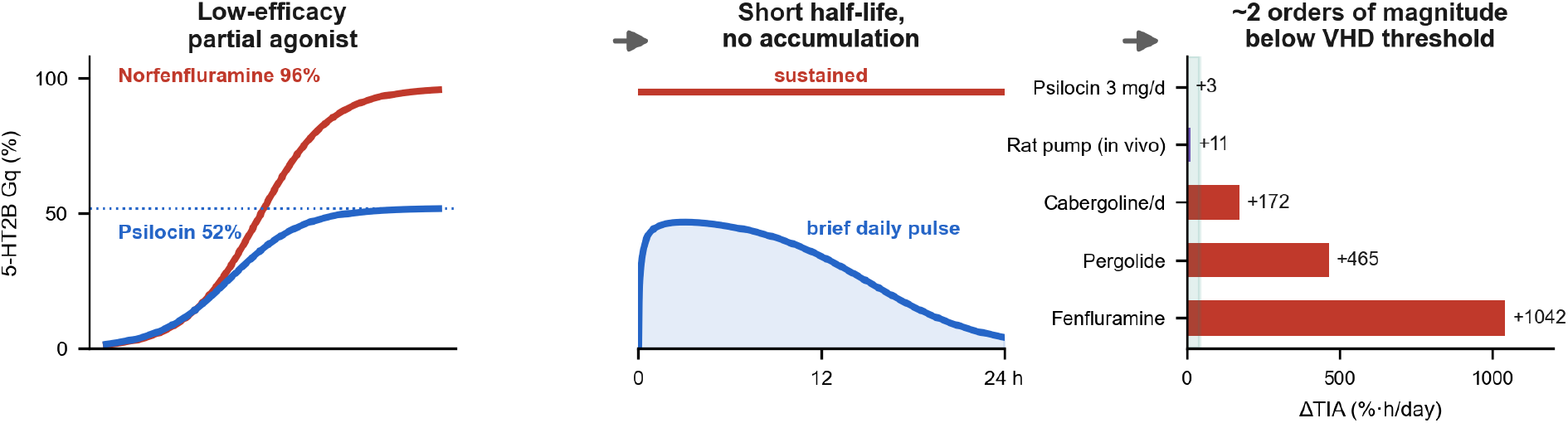

Three key determinants of cardiac safety margins for repeated low-dose psilocybin are shown. Psilocin is a low-efficacy partial agonist at 5-HT2B (ceiling ≈52% vs 96% for norfenfluramine; left). Its short half-life yields a brief daily pulse of receptor engagement rather than a sustained plateau (center). The resulting integrated 5-HT2B signal (ΔTIA) at 3 mg daily lies roughly two orders of magnitude below valvulopathic exposures, and continuous in vivo exposure produced no valvulopathy (right).

## 1. INTRODUCTION

The therapeutic development of psilocybin has proceeded almost entirely through single high (“macrodose”) administrations of 10-25 mg delivered under intensive psychological support. Although efficacious in trials for treatment-resistant depression, this paradigm is difficult to scale because it demands specialized settings, trained personnel, prolonged monitoring, and, prospectively, burdensome risk-management infrastructure. An alternative is the repeated administration of low, sub-perceptual doses (e.g., 2-5 mg) on a self-administered, outpatient basis. This sub-hallucinogenic paradigm has seen substantial community uptake with an estimated 11 million low-dose psilocybin use days in the United States in 2025 supported by growing preclinical rationale in models of motivation, addiction, attention and mood [1-7]. If safe and effective, such a regimen would be far more accessible than high dose interventions. Its central pharmacological distinction from high dose therapy, however, is the exposure pattern. Rather than isolated high peaks separated by weeks or months in high dose approaches, low dose methods of treatment produce frequent, repeated receptor engagement over months or years. Differences in these exposure profiles raise critical safety questions remaining to be resolved.

A risk of chronic serotonergic agonist use is drug-induced valvular heart disease. A well-defined class of agents such as the fenfluramine anorectics, the ergot-derived dopamine agonists pergolide and cabergoline, and methysergide have been demonstrated to produce valvular fibrosis. The shared mechanism for this off-target side effect is agonism at the 5-HT2B receptor expressed on cardiac valve interstitial cells [8-11]. Excessive activation of this G_q/11_-coupled receptor drives mitogenic, transforming growth factor-β1 (TGF-β1) dependent signaling in a manner sufficient to increase cardiac valvular fibrosis risks [12-14]. Indeed, the clinical precedents are unambiguous where fenfluramine– phentermine produced a characteristic valvulopathy first described in a case series of 24 users, all with valvular regurgitation [15]. Moreover, the ergot dopamine agonists pergolide and cabergoline carry a cumulative-dose– dependent risk of valvular regurgitation [16]. Endogenously, the same biology underlies carcinoid heart disease, in which chronically elevated circulating serotonin correlates with valvular damage [17].

Psilocin binds with nanomolar affinity acting as a partial agonist on 5-HT2B receptors, so questioning its effects on cardiopathogenesis is legitimate and cannot be dismissed by assertion [18, 19]. Regulatory guidance suggests sponsors of psychedelic drugs trials should characterize functional 5-HT2B activity and, for agonists intended for chronic use, to conduct heart-valve histopathology in repeat-dose toxicology and prospective echocardiography clinical trials [20]. Scientific discourse on the risks for cardiac valvular fibrosis associated with psilocybin use have largely relied on hypotheses supported by serotonin receptor binding affinities. This incomplete profiling approach elevates risk perception since other pharmacological factors are not considered. Here based on established pharmacology, we propose that functional efficacy and exposure duration, not affinity, are major contributors to valvular fibrosis risk. To illustrate this perspective, we introduce a simple, transparent exposure–response model that estimates 5-HT2B signaling produced by a candidate low-dose psilocybin dosing regimen. We calibrated this model using other agonist exposures with established outcomes. We also provide preliminary *in vivo* cardiac histopathology data as corroboration of the model. We are explicit about the limitations of evidence supporting the model, as well as further work required to validate it. This initial analytical framework is intended to help establish new models for characterizing the risk of repeated, low-dose psilocybin clinical research programs, as well as those using other non-selective agonists that have an affinity for 5-HT2B receptors.

## 2. METHODS

### 2.1. Receptor pharmacology parameters

Fractional 5-HT2B G_q_ activation was modeled by the Hill relation (**Eq. 1**), with potency (EC_50_) and efficacy (E_max_, expressed as a percentage of the serotonin maximum) taken from a single matched G-protein dissociation (BRET) assay at the human 5-HT2B receptor [21].

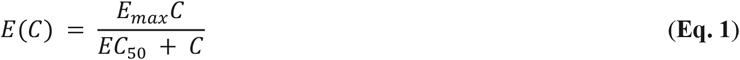

Here E(C) is the fractional G_q_ activation at agonist concentration C, E_max_ the efficacy ceiling of the ligand and EC_50_ its half-maximal concentration (Hill coefficient 1). The matched-assay values for serotonin, psilocin, LSD and norfenfluramine are shown in **Table 1** below. Pergolide and cabergoline have been shown to be potent, high-efficacy 5-HT2B agonists in a matched multi-drug functional panel (E_max_ spanning ≈60–108% of the serotonin maximum; [22]). These drugs were modeled with representative mid-range parameters reflecting their high 5-HT2B affinity and efficacy and are flagged as lower confidence (**Table 1**).

**TABLE 1.** Functional 5-HT2B Gq Parameters Used in the Exposure–Response Model.

| Ligand | EC<sub>50</sub> (nM) | E<sub>max</sub> (% of 5-HT max) | Confidence | Source |
| --- | --- | --- | --- | --- |
| Serotonin (5-HT) | 0.95 | 100 | High | [21] |
| Psilocin | 3.10 | 51.8 | High | [21] |
| LSD | 0.88 | 62.1 | High | [21] |
| Norfenfluramine | 6.78 | 96.2 | High | [21] |
| Pergolide | 7.1 | 90 | Low | [22] |
| Cabergoline | 1.2 | 75 | Low | [22] |

### 2.2 Pharmacokinetic model

Plasma psilocin was described by a one-compartment model with first-order absorption and elimination (**Eq. 2**), scaled to the observed peak.

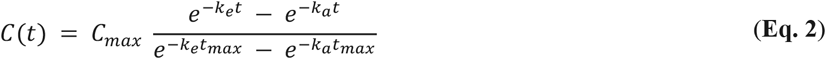

where k_e_ and k_a_ are the first-order elimination and absorption rate constants and t_max_ the time of peak concentration. The denominator normalizes the profile so that C(t_max_) = C_max_. Parameters for oral psilocybin were fitted to data from our Phase 1 single-ascending-dose study (n = 56; 0.5–4.0 mg; [23]), giving a dose-proportional C_max_ of ≈3.7 ng/mL (18.0 nM) at 3 mg, t_max_ ≈ 1 h, and terminal half-life ≈ 2.5 h. Steady-state profiles were obtained by superposing successive doses at the specified interval until convergence (≥10 half-lives). Comparator peak plasma concentrations and absorption/elimination parameters were taken from the compilation of Rouaud et al. (2024), sub-hallucinogenic LSD from Holze et al., (2021), d-norfenfluramine from Richards et al., (1989), cabergoline from Andreotti et al., (1995) and pergolide (2.73 ng/mL) from Blin (2003) as shown in **Table 2** [18, 24-27]. The LSD value of 0.28 ng/mL corresponds to a 10 µg daily dose where a 20 µg dose would be ≈0.50 ng/mL [26]. For pergolide, a dedicated multiple-dose study has reported lower steady-state concentrations (≈0.1–1.2 ng/mL) [28]. As a sensitivity check we note that adopting these lower values reduces the pergolide score without altering the psilocin margin. Concentrations were converted to molar units using each compound’s molecular weight (psilocin 204.27 g/mol).

**TABLE 2.** Pharmacokinetic Inputs Used to Model 5-HT2B Agonist Exposure.

| Drug (regimen) | C<sub>max</sub> (ng/mL) | t<sub>max</sub> (h) | t<sub>1/2</sub> (h) | Interval (h) | Source |
| --- | --- | --- | --- | --- | --- |
| Psilocin (3 mg daily) | 3.7 | 1.0 | 2.5 | 24 | [23] |
| LSD (10 μg daily) | 0.28 | 1.5 | 3.6 | 24 | [18, 26] |
| d-Norfenfluramine (fenfluramine 30 mg BID) | 21 | 8.0 | 30 | 12 | [18, 27] |
| Pergolide (1 mg daily) | 2.73 | 2.0 | 27 | 24 | [18, 25] |
| Cabergoline (3 mg daily) | 0.12 | 2.5 | 65 | 24 | [18, 24] |
| Cabergoline (1 mg weekly) | 0.04 | 2.5 | 65 | 168 | [18, 24] |

### 2.3 Exposure metric and competitive model

Because valve interstitial cells reside in endogenous serotonin, an exogenous agonist was modeled as competing with 5-HT at the receptor rather than acting on a naive receptor. The net fractional activation therefore follows a two-ligand competitive form (**Eq. 3**):

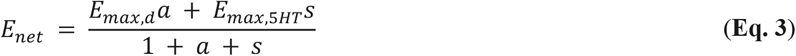

where a = C_d_/EC_50,d_ and s = C_5HT_/EC_50,5HT_ are the normalized drives of drug and serotonin, each contributing its efficacy weighted by occupancy while the shared denominator (1 + a + s) captures competition for the receptor (all symbols defined in **Table 3**). Free valvular serotonin was set to a nominal 1 nM (varied in sensitivity analysis). This value reflects the free, platelet-poor pool that valve interstitial cells sense, which is low-nanomolar because the great majority of circulating 5-HT is sequestered in platelets (whole-blood and platelet-rich plasma concentrations are far higher; Bhattacharyya et al., 2013). The choice is deliberately conservative for the candidate: because raising the assumed baseline increases endogenous competition and therefore lowers the computed increment for every exogenous agonist, any higher, more physiologic free-serotonin value would only reduce the 3 mg daily score below +3, never raise it. The primary metric was the increment in net drive above the endogenous baseline integrated over one 24 h interval at steady state (**Eq. 4**):

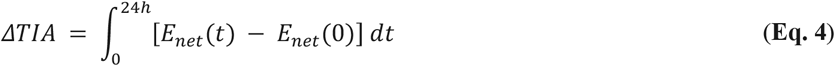

expressed in percent-hours per day, where E_net_(0) is the baseline activation with no drug present. A modeling of endogenous competition is required because a naive occupancy model saturates and cannot distinguish healthy serotonin tone from carcinoid heart disease. The duty cycle (hours per day with increment ≥ 10 percentage points) was reported alongside ΔTIA because the fibrotic program depends on sustained signaling. A conservative single-ligand variant (no endogenous competition) was computed as a worst case. Endogenous-serotonin reference states (median plasma 5-HT: healthy 9 nM, carcinoid syndrome 155 nM, carcinoid heart disease 325 nM; [29]) were scored on the same increment scale. Calculations were performed in Python (NumPy) by numerical integration; the model is deterministic and fully specified by the parameters above (**Table 3**).

**TABLE 3.** Model Quantities and Definitions for the 5-HT2B Exposure–Response Framework.

| Quantity | Definition | Units / value |
| --- | --- | --- |
| $E(C)$ | Fractional 5-HT2B Gq activation at agonist concentration C (Hill relation, Eq. 1). | % of 5-HT max |
| $E_{max}$ | Efficacy ceiling — the maximal activation a ligand can produce (partial agonism if below 100%). | % of 5-HT max |
| $EC_{50}$ | Potency — agonist concentration giving half-maximal activation. | nM |
| $C$ | Ligand concentration at the receptor. | nM |
| $C_{max}$ | Peak plasma concentration (per drug; Table 2). | ng/mL |
| $k_e, k_a$ | First-order elimination and absorption rate constants ($k_e = \ln 2 / t_{1/2}$). | $h^{-1}$ |
| $t_{max}$ | Time of peak plasma concentration. | h |
| $E_{net}$ | Net activation with drug and endogenous 5-HT competing at the receptor (Eq. 3). | % of 5-HT max |
| $E_{max,d}, E_{max,5HT}$ | Efficacy of the exogenous drug d and of serotonin ($E_{max,5HT} = 100\%$). | % |
| $a$ | Normalized drug drive, $a = C_d / EC_{50,d}$. | dimensionless |
| $s$ | Normalized serotonin drive, $s = C_{5HT} / EC_{50,5HT}$. | dimensionless |
| $C_d$ | Free drug (e.g., psilocin) concentration at the valve. | nM |
| $C_{5HT}$ | Free (platelet-poor) valvular serotonin concentration. | nM; nominal 1 |
| $\Delta TIA$ | Time-integrated increment in Gq drive above endogenous tone over one 24 h interval at steady state (Eq. 4). | %·h/day |
| Duty cycle | Hours per day with the increment at least 10 percentage points above baseline. | h/day |

### 2.4 Cardiac histopathology

The cardiac observations summarized in Section 2.5 were not generated for the present work. They were derived from a sponsored preclinical rat study conducted by a contract research organization for a behavioral (5-choice serial reaction time task) endpoint (Study Report IVS209-20064-RO; [30]). In that study, male Long-Evans rats (Charles River) received continuous subcutaneous psilocin from Alzet osmotic minipumps (model 2ML2) primed with saline (N = 16) or with psilocybin 5 mg/mL (N = 24) for twelve days, producing a steady plasma psilocin of ≈8–10 ng/mL (LC-MS/MS on days 4, 8 and 12). All animal procedures were performed in accordance with the principles of the Canadian Council on Animal Care (CCAC). Hearts from 6 vehicle and 6 psilocybin pump-implanted rats, randomly selected from the study, were subsequently provided to an independent pathology laboratory, where a certified histopathologist masked to treatment (Dr. Susan Camilleri, TCP Pathology Core, Toronto Centre for Phenogenomics) examined longitudinal hematoxylin-and-eosin sections of all four heart valves and of the myocardium, endocardium and epicardium for degeneration, thickening and other abnormalities.

## 3 RESULTS

### 3.1 Binding affinity does not predict valvular outcomes

Drug-induced valvulopathy is caused by over-signaling of proliferative cellular molecular processes. Thus, the pharmacological quantity that should predict it is functional 5-HT2B agonism, not the tightness of binding alone. Consistent with this, a binding-affinity rule can misclassify agonists with established human outcomes (**Table 4**). Norfenfluramine (the valvulopathic metabolite of fenfluramine) and MDMA both cause valvular heart disease (VHD) despite weak 5-HT2B binding (K_i_ ≈ 52 and 500 nM). This is because they produce robust receptor signaling under sustained exposure. Conversely, several antipsychotics and the ergoline lisuride bind 5-HT2B with sub-nanomolar affinity yet carry no valvular risk, because they are antagonists or inverse agonists with negligible intrinsic efficacy [18]. A parallel functional-activity profiling study established that the drugs causing VHD are potent, high-efficacy 5-HT2B agonists and functional agonism, not binding affinity alone, is their common property [22]. This is also a functional axis on which regulatory guidance is framed [20]. **Table 4** provides K_i_ values from a valvular pathogenic subset of compounds [18] in the NIMH Psychoactive Drug Screening Program (PDSP) database. A binding-affinity screen flagging agents whose 5-HT2B K_i_ lies within roughly an order of magnitude of the K_i_ for serotonin [18, 31] as an approximate low-nanomolar cutoff is a reasonable trigger, not a risk metric, as shown illustrated in **Table 4**.

**TABLE 4.** Discordance Between 5-HT2B Binding Affinity and Valvular Outcomes.

| Drug | 5-HT<sub>2B</sub> $K_i$ (nM) | Affinity rule | Human outcome | Pharmacology |
| --- | --- | --- | --- | --- |
| Norfenfluramine | 52 | Low risk (>15) | Causes VHD | Near-full agonist |
| MDMA | 500 | Low risk (>15) | Causes VHD | 5-HT releaser + agonist |
| Lisuride | 2.9 | High risk (<15) | No VHD | Antagonist |
| Aripiprazole | 0.36 | High risk (<15) | No VHD | Inverse agonist |
| Psilocin | 4.7 | High risk (<15) | Low-efficacy partial agonist | See sections 3.2–3.4 |

### 3.2 Psilocin is a low-efficacy partial 5-HT2B agonist

In a matched G-protein dissociation assay at the human receptor, psilocin activates 5-HT2B with an efficacy ceiling of 51.8% of the serotonin maximum, compared with 96.2% for norfenfluramine and 62.1% for LSD (**Figure 1**) [21]. Four independent assays demonstrate psilocin’s 5-HT2B efficacy to be concordantly in a 38–52% range (reviewed by [19]). Because a partial agonist cannot exceed its ceiling at any concentration, psilocin is intrinsically incapable of driving valvular G_q_ signaling beyond roughly half of that which a full agonist or endogenous serotonin can produce. This efficacy gap, not differences in binding affinities, is the pharmacological basis of the safety margin quantified below.

**FIGURE 1.**
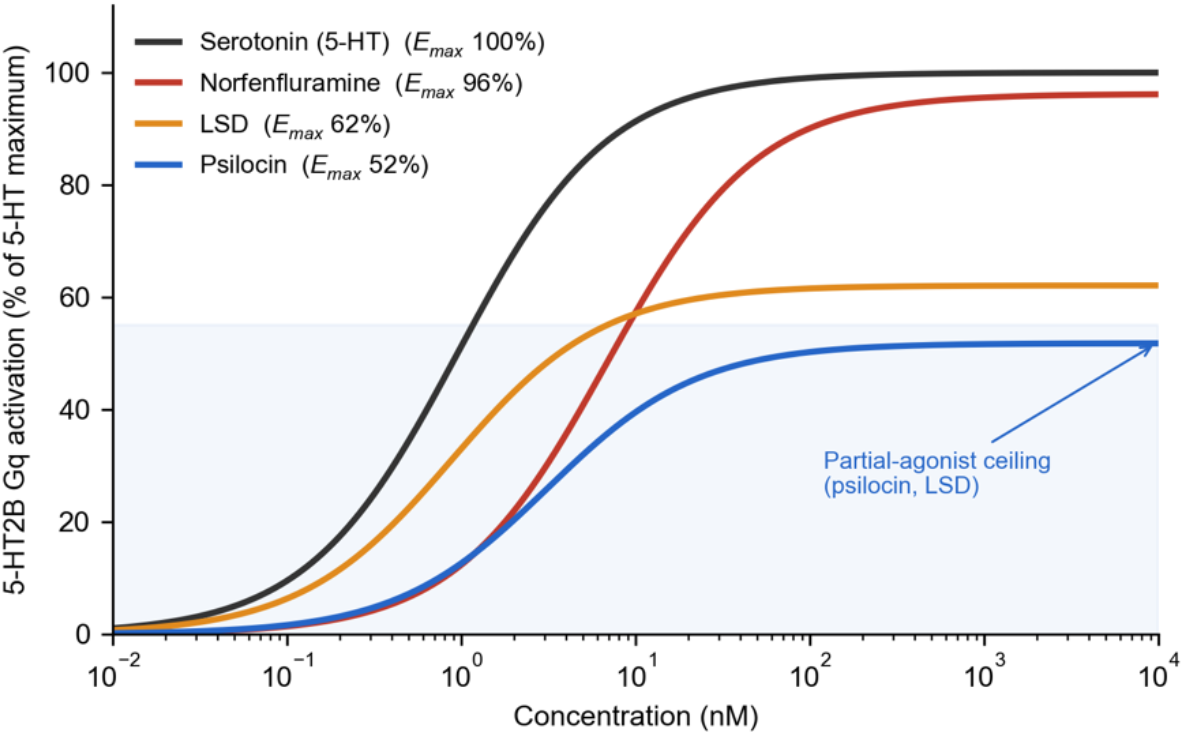
Human 5-HT2B Gq concentration–response relationships. The line plots illustrate a plateau height corresponding to the maximal signaling each ligand can produce at any concentration. Psilocin (E_max_ 52%) and LSD (E_max_ 62%) are partial agonists, whereas the valvulopathic metabolite norfenfluramine (E_max_ 96%) approaches the serotonin maximum. Curves computed from matched-assay potency and efficacy values [21].

### 3.3 Exposure–Response Modeling Places 3 mg Daily Psilocybin Below Valvulopathic Exposures

To model and quantify valvulopathic risk beyond binding affinities, we calculated the time-integrated increase in 5-HT2B G_q_ activation above endogenous serotonergic tone across a 24 h steady-state dosing interval. We define this metric as ΔTIA, expressed in percent-hours per day (see Section 3 Methods). Plasma psilocin is described by a one-compartment model fitted to data from our Phase 1 single-ascending-dose study in healthy adults [23], which gives a dose-proportional peak concentration of ≈3.7 ng/mL (≈18 nM) at 3 mg with a terminal half-life of ≈2.5 h. The fractional 5-HT2B activation is computed from the Hill equation using the matched-assay parameters (see Section 2.2 above), with the drug modeled as competing with endogenous serotonin at the receptor binding site. We calibrated the model against exposures of known outcomes. The model yields a discriminating scale with no misclassifications across the molecules examined (**Table 5 and Figure 2**). Every exposure modeled that causes VHD had scores ΔTIA ≥ +172 %·h/day. For example, norfenfluramine (+1042), pergolide (+465), daily high-dose cabergoline (+172), and, as an endogenous reference, carcinoid heart disease (+1108) fell into this category. Exposures modeled that not been shown to cause VHD scored ≤ +28 with examples being low-dose weekly cabergoline (+21), daily sub-hallucinogenic 10 µg LSD (+28), and as baseline reference normal serotonin tone (+0).

**TABLE 5.** Calibration of 5-HT2B Exposure–Response Metrics Against Valvular Outcomes.

| Exposure | ΔTIA | Sustained drive (h/day) | Human VHD? |
| --- | --- | --- | --- |
| Endogenous 5-HT - carcinoid heart disease | +1108 | 24 | Yes |
| d-Norfenfluramine (fenfluramine 30 mg BID) | +1042 | 24 | Yes |
| Pergolide (1 mg daily) | +465 | 24 | Yes |
| Cabergoline (3 mg daily, Parkinson's) | +172 | 24 | Yes |
| LSD (10 μg daily) | +28 | 0 | No |
| Cabergoline (1 mg weekly, prolactinoma) | +21 | 0 | No |
| Rat pump - psilocin 9.2 ng/mL continuous ×12 d | +11 | 0 | No (*in vivo* rat) |
| **Psilocin - 3 mg psilocybin daily (candidate)** | **+3** | **0** | — |
| Endogenous 5-HT - healthy (9 nM) | +0 | 0 | No |

Rows sorted by ΔTIA. Psilocin, LSD, norfenfluramine and serotonin use matched-assay efficacies [21]; pergolide and cabergoline are potent, high-efficacy 5-HT2B agonists (functional $E_{\max}$ spanning ≈ 60–108% of the 5-HT maximum across assays; [22]) and are modeled with representative mid-range parameters (lower confidence). Pergolide is scored at its compiled peak plasma concentration (2.73 ng/mL; [18, 25]); at the lower steady-state concentrations reported by a dedicated multiple-dose study ( ≈ 0.1–1.2 ng/mL; [28]) its score falls toward the +172 threshold ( ΔTIA ≈ +145 to +236 across that range) but remains far above the 3 mg psilocybin candidate (+3), so the qualitative separation is unchanged. Serotonin calibration anchors are median plasma 5-HT values (healthy ≈ 9 nM; carcinoid syndrome ≈ 155 nM; carcinoid heart disease ≈ 325 nM; [29]).

**FIGURE 2.**
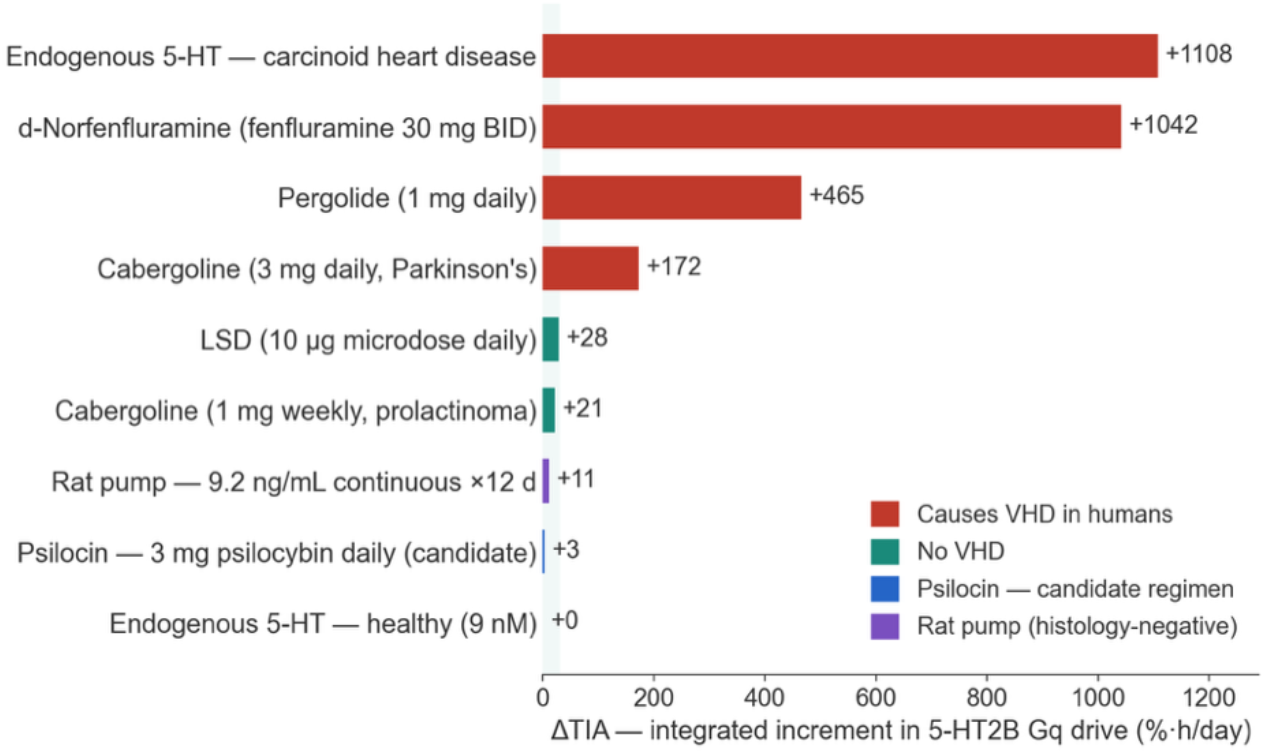
Integrated 5-HT2B signaling (ΔTIA) across exposures with known valvular outcomes. The color-coded horizontal bars illustrate the integrated 5-HT2B drive in percent-hours per day (ΔTIA) where: red = human VHD; green = no VHD; blue = 3 mg daily psilocin; and purple = *in vivo* rat osmotic-pump results. Psilocin at doses shown falls in the safe band, below valvulopathic exposure risk thresholds. Shading marks the empirical no-VHD zone.

A 3 mg daily psilocybin regimen scores a ΔTIA of +3, below even the negative controls and roughly two orders of magnitude below the weakest exposure that has caused human valvulopathy. The LSD comparator is modeled at a daily sub-hallucinogenic 10 µg dose (peak 0.28 ng/mL; [26]), which scores +28. This dose of LSD together with weekly cabergoline (+21) set the upper edge of the calibrated no-VHD band. A 20 µg daily LSD dose would raise the peak to ≈0.50 ng/mL and the ΔTIA score to +44. Although this exceeds the calibrated ceiling, it is supported by direct safety data. Chronic (4- and 8-weeks; 5 days/week) sub-hallucinogenic LSD dosing (0.01 and 0.03 mg/kg i.p.) failed to cause ventricular or valvular remodeling by echocardiography in mice, whereas the positive controls d-fenfluramine (10 mg/kg, i.p.) and exogenous serotonin (40 mg/kg, i.p.) did [21]. Eight weeks of up to 20 µg LSD twice weekly produced no echocardiographic valvulopathy in patients with major depressive disorder [32]. Both the 10 and 20 µg exposures therefore remain far below the +172 valvulopathic threshold and are concordant with the observed absence of valve disease, while the daily 3 mg psilocybin candidate (ΔTIA +3) sits an order of magnitude lower still.

Cabergoline is instructive: an identical receptor pharmacology carries a cumulative-dose–dependent valvular risk at the high daily doses used in Parkinson’s disease [16] but not at the low intermittent (≤ weekly) doses used for hyperprolactinemia, and the model reproduces this separation on exposure alone (daily +172 versus weekly +21)— illustrating that exposure pattern, not affinity, governs the outcome. Under a deliberately conservative variant that ignores endogenous competition, the integrated-signal margin of 3 mg daily over fenfluramine remains ≈5.5-fold.

### 3.4 Psilocin Produces Pulsatile Rather Than Sustained 5-HT2B Engagement

Beyond the efficacy ceiling, drug exposure duration independently favors safety. Because the half-life of sub-hallucinogenic, orally administered psilocin (≈ 2.5 h; [23]) is short relative to a 24 h dosing interval, superposition of daily doses produces no accumulation (steady-state/single-dose ratio 1.00) and a return to near-zero plasma concentrations for approximately twenty hours each day (**Figure 3A**). Consequently, lower sub-hallucinogenic doses once per day engage 5-HT2B as a brief daily pulse that decays to baseline, whereas the valvulopathic drugs probed maintain a sustained plateau of receptor activation (**Figure 3B**). Our exposure-response model attributes zero hours per day of sustained, elevated 5-HT2B drive to 3 mg daily. This contrasts with persistent 5-HT2B activity elicited by fenfluramine (**Table 5**). Because the fibrotic transcriptional program requires continuous rather than intermittent stimulation [13], this continuous *vs* intermittent distinction (instead of any single potency ratio) highlights potential safety margins.

**FIGURE 3.**
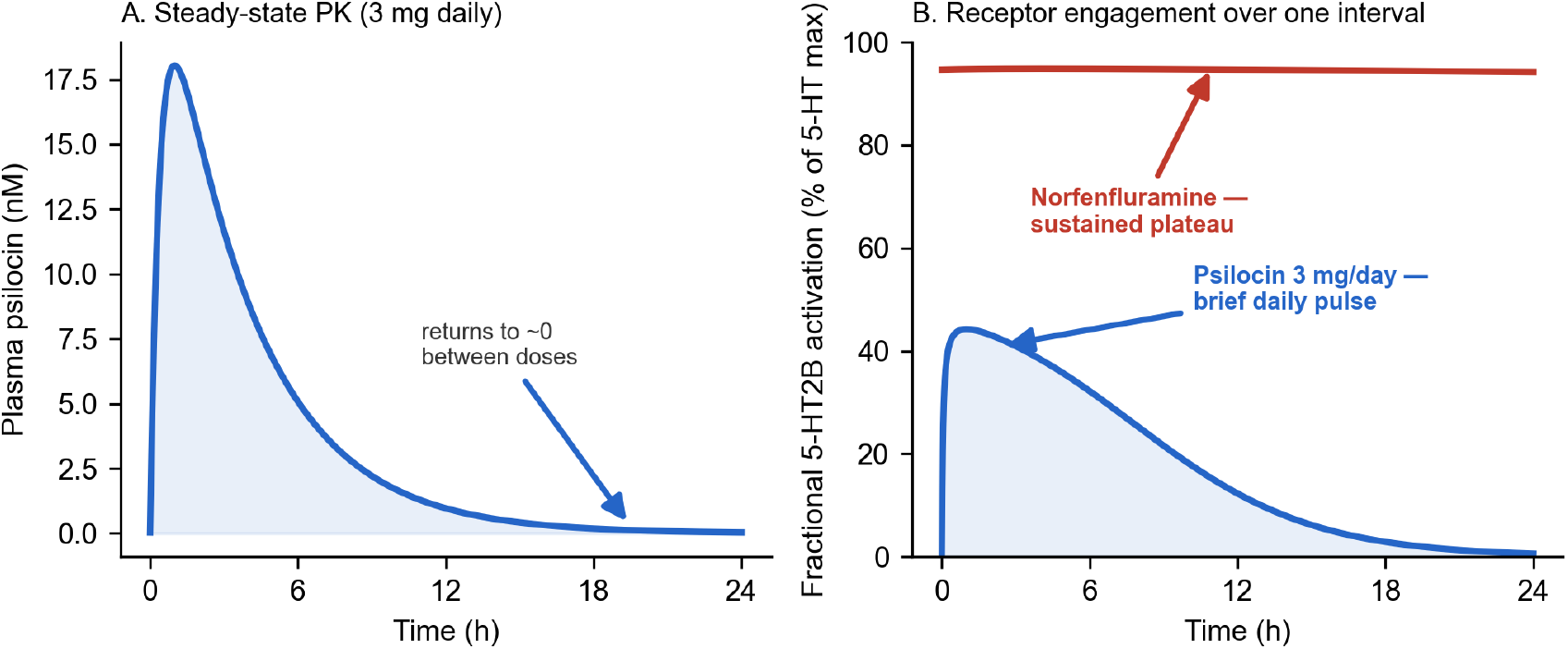
Pulsatile and continuous 5-HT2B engagement highlight safety margins for sub-hallucinogenic psilocybin. (A) The psilocin PK curve for 3 mg daily psilocybin modeled from our prior observations illustrates no accumulation and complete return to near-zero between doses. (B) The plot illustrates the fractional and sustained activation of 5-HT2B over a 24-hour period. Psilocin (blue) engages the receptor as a brief daily pulse (A) that decays to baseline, whereas norfenfluramine (red) maintains a sustained plateau at nearly complete (100%) engagement.

### 3.5 Preclinical Cardiac Histopathology Supports the Exposure–Response Model

We compared the exposure-response model against cardiac tissue that was already available from one of our preclinical programs, independent of the present analysis. In that study conducted to examine a behavioral endpoint (Study Report IVS209-20064-RO; [30]), male Long-Evans rats received continuous psilocin from a subcutaneous osmotic minipump (Alzet model 2ML2) for 12 days. The serial pharmacokinetics confirmed stable, continuous plasma levels of 10.0, 8.7 and 8.9 ng/mL on days 4, 8 and 12 with a mean of 9.2 ng/mL. This daily, continuous exposure is 2.4-fold above the projected peak of a 3 mg daily dose in humans illustrated (**Figures 2 and 4**). Because this profile happens to reproduce the sustained agonism relevant to valvular safety, hearts from 6 vehicle-pump and 6 psilocybin-pump rats, randomly selected from the study, were provided to an independent laboratory and examined using hematoxylin-and-eosin sections through the valves by a pathologist blinded to treatment (S. Camilleri, TCP Pathology Core). We include this observation as external corroboration of the exposure-response model, not as a primary observation of the present paper.

**FIGURE 4.**
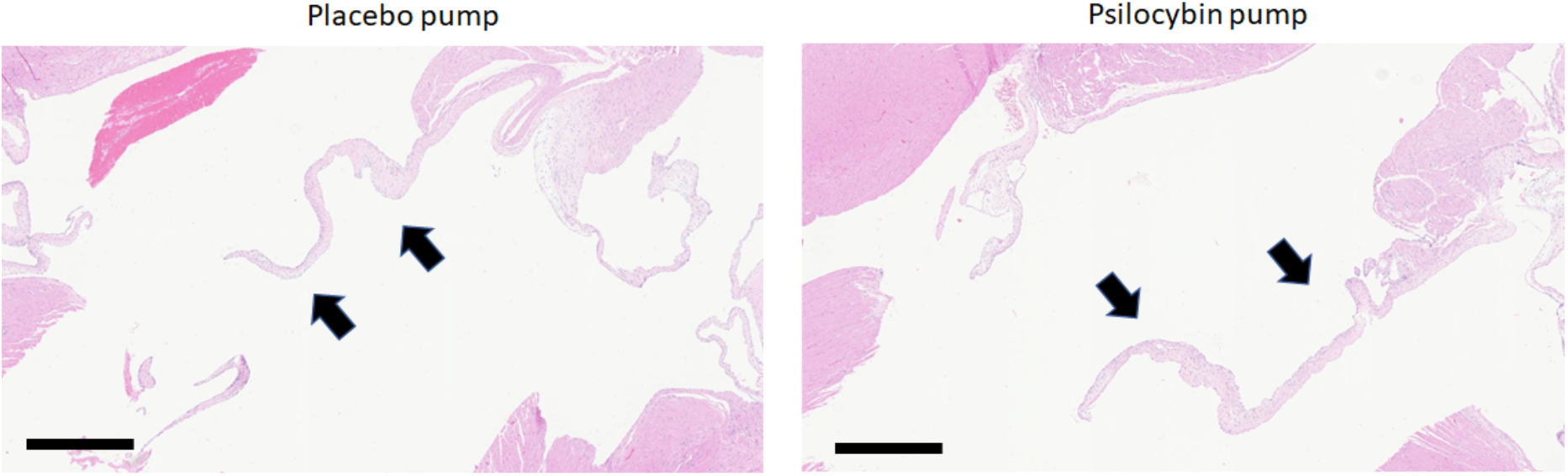
Cardiac histopathology corroborates the exposure-response model. Photomicrograph of the aortic valve (arrows) from a rat implanted with an Alzet minipump primed with saline (placebo pump, left) or with psilocybin 5 mg/mL (right) producing continuous psilocin plasma levels (≈8–10 ng/mL) over 12 treatment days. No lesions were observed in the myocardium, endocardium or epicardium. No abnormalities were observed in the aortic or mitral valves. Overall, all cardiac tissues appeared normal across the 6 placebo-pump and 6 psilocybin-pump rats examined (representative sections shown). Cardiac histopathology was conducted in a blinded fashion by a certified histopathologist (Dr. Susan Camilleri, Toronto Centre for Phenogenomics, TCP Pathology Core), on tissue from a rat behavior study [30]. Scale bar = 0.5 mm.

No lesions were observed in the myocardium, endocardium, epicardium, or the aortic or mitral valves of hearts from psilocin-exposed rats and they were indistinguishable from vehicle controls (**Figure 4**). The independent, blinded report concluded that all cardiac tissues appeared normal, with no histological evidence of valvulopathy (S. Camilleri, TCP Pathology Core; supplemental to [30]). This rat model has been demonstrated as valid and sensitive model of serotonergic valvulopathy and has been used to serotonin-induced aortic-valve regurgitation by echocardiograph [33]. Other independent studies show chronic serotonin or serotonergic drugs produce valvular thickening and regurgitation in the rat with histological confirmation, in a dose-dependent and reversible manner [34-36]. Companion behavioral measures showed no serotonin-syndrome or cardinal 5-HT2A signs and no change in locomotor activity across these exposures (Study Report IVS209-20064-RO; [30]).

While the preliminary corroboration of the exposure-response model is encouraging, appropriate cautions, proactive clinical monitoring, and transparent reporting remains needed. In the positive-control models cited above, serotonergic VHD develops over sustained exposures of several weeks to months rather than days. The twelve-day rat study therefore establishes acute-to-subacute cardiac tolerability at a supra-clinical continuous exposure while confirming the model’s short-term prediction. However, the model cannot exclude a slowly developing fibrosis stemming from chronic (multi-month) exposure. Therefore, these findings are best taken as an early-timepoint observation concordant with the exposure–response model and is not as a substitute for chronic repeat-dose toxicology. Placed on the model’s scale, the continuous pump exposure scores ΔTIA +11 within the safe band while the preliminary histopathology agrees (**Table 5 and Figure 2**).

## 4 DISCUSSION

Three independent lines of evidence converge through observations. First, from established pharmacology, valvular risk tracks functional efficacy and exposure duration rather than binding affinity, and psilocin is a high-affinity but low-efficacy partial agonist at 5-HT2B. Second, an exposure–response model built on matched-assay pharmacology and Phase 1 pharmacokinetics, calibrated without error against seven clinical controls, places a 3 mg daily regimen roughly two orders of magnitude below the weakest valvulopathic exposure. This is driven jointly by the partial-agonist ceiling and a short half-life of psilocin that precludes accumulation. Third, continuous *in vivo* exposure above the projected 3 mg daily peak in for 12 days produced no valvulopathy in a rat model with demonstrated sensitivity to serotonergic valve disease. These data suggest daily, low-dose psilocybin lacks the features of a drug poised to cause valvular fibrosis.

Emerging human data are consistent with this quantitative model. In the first clinical trial to bracket repeated psychedelic dosing with transthoracic echocardiography, eight weeks of twice-weekly LSD microdosing in patients with major depression produced no echocardiographic evidence of valvulopathy [32]. LSD is a more efficacious 5-HT2B agonist than psilocin (62.1% versus 51.8% in the matched assay data used here), making this a stringent, albeit early, human precedent. At the population scale, an analysis of the All of Us cohort found that lifetime hallucinogen use was, in crude comparison, associated with a lower prevalence of valvular heart disease (3.6% versus 4.7%) [37]. Interestingly, only after adjustment and remodeling for demographic and clinical confounders did the association reverse to a modest, marginally significant increase (adjusted odds ratio 1.08, 95% CI 1.01–1.55) [37]. Such a small, adjustment-dependent signal derived from undated lifetime exposure of unknown dose, unknown formulation and frequency, and confounded by co-used substances including MDMA cannot establish a causal, dose-related effect of controlled low-dose psilocin. It does however highlight the plausible magnitude of any real-world risk as small. The convergence of a null mechanistic model, a null short-term animal study, a null human echocardiographic trial, and at most a marginal epidemiological association is reassuring. Despite these encouraging data, they are not a substitute for prospective monitoring in the specific dosing regimen under development.

Several limitations temper our conclusions and define the necessary work ahead. Psilocin’s partial agonism is defined here on the G_q_ pathway, on which it is a low-efficacy partial agonist (≈38–52% of the serotonin maximum across matched assays; [21, 38]). At 5-HT2B, however, psilocin recruits β-arrestin2 with substantially higher efficacy (≈76– 84%) than it activates the G_q_/calcium arm. Restated, psilocin is arrestin-preferring rather than balanced at this receptor [38]. Future studies need to resolve whether β-arrestin activation by psilocin is fibrogenic. This should also be addressed through matched G_q_/β-arrestin profiling, ideally in a multi-drug functional panel [22, 39]. Another limitation is the animal evidence corroborating our model only had an exposure span of 12 days, whereas the intended clinical use spans months to years. Although psilocin shows no accumulation and lacks the serotonin-releasing mechanism that plausibly underlies MDMA-associated valvulopathy, the effects of chronic pulsatile 5-HT2B agonism over years has not been investigated. Thus, no short-term study can exclude a slowly emerging side effect. Finally, the model depends on parameters that carry uncertainty. For instance, uncertain parameters include psilocin’s unbound fraction, the free serotonin concentration at the heart valves, and use of literature-derived comparator efficacies which we have made explicit. Across the proposed daily psilocybin dose range studied, the qualitative conclusion is stable, but the precise margin is not a single fixed value.

Based on these observations, prudence is warranted. However, the appropriate course is continued clinical investigation under quantified vigilance rather than dismissal of the concern or inaction driven by uncertainty, especially considering emerging preclinical, population, and clinical data regarding safety and risks [4, 19, 21, 32, 37]. Consistent with regulatory expectations for chronically administered 5-HT2B agonists [20], clinical trials of repeated low-dose psilocybin should include prospective echocardiography from baseline through periodic follow-up to assess valve structure, valve function, and pulmonary-artery pressures. Trials should also exclude participants with pre-existing valvulopathy or pulmonary hypertension. Clinical programs may further consider intermittent dosing schedules to reduce cumulative exposure, an approach supported by evidence that serotonin-induced valvular changes in rats are dose-dependent and reversible after withdrawal [35]. This strategy mirrors the monitoring framework used for an approved 5-HT2B agonist and reframes the objective from proving zero risk to characterizing and managing a quantified risk. On the present evidence, that objective appears achievable, and repeated low-dose psilocybin merits continued, carefully monitored clinical investigation for neuropsychiatric disorders.

## DECLRATIONS

## Author Contributions

W.J.T. conceived the analysis and drafted the manuscript. W.J.T., M.B.M., and E.S. contributed to the pharmacological and regulatory framework, interpreted the data and revised the manuscript. All authors approved the final version. The *in vivo* rat study was directed by G. A. Higgins (InterVivo Solutions Inc.); the cardiac histopathology reproduced in Figure 4 was performed independently and blinded by Dr. Susan Camilleri (TCP Pathology Core, Toronto Centre for Phenogenomics). These contributors are acknowledged rather than listed as authors of this perspective.

## Acknowledgments

We thank InterVivo Solutions Inc. for conducting the sponsored rat study (IVS209-20064-RO), and the Toronto Centre for Phenogenomics (TCP) Pathology Core, and Dr. Susan Camilleri, for the independent blinded cardiac histopathology.

## Animal Ethics

The rat study from which the cardiac tissue derives (Sellers and Higgins, 2020) was conducted following all institutional and national guidelines for the care and use of laboratory animals; all animal use procedures were performed in accordance with the principles of the Canadian Council on Animal Care (CCAC).

## Data Availability

Model parameters and equations are given in full in Section 3. Pharmacological and clinical values derive from the cited literature. The Phase 1 pharmacokinetic dataset (in preparation) and the sponsored rat study report (Sellers and Higgins, 2020; Study Report IVS209-20064-RO), together with its supplemental cardiac histopathology, are available from the corresponding author on reasonable request.

